# Antimycin A, but not antimycin A3 or myxothiazol, directly suppresses photosystem II activity

**DOI:** 10.1101/2025.09.11.675326

**Authors:** Ko Imaizumi, Kentaro Ifuku

## Abstract

Antimycin A (AA) is a widely used inhibitor to study photosynthesis and respiration. In photosynthesis, it is commonly used to inhibit a cyclic electron flow (CEF) pathway but has also been reported to affect photosystem II (PSII), which is not involved in CEF. Although concerns have been raised about AA’s specificity, its impact on PSII activity remains unclear. AA3 was recently proposed as a more specific inhibitor of the same CEF pathway. In the mitochondrial respiratory chain, AA inhibits complex III, similar to myxothiazol. Here, we investigated the direct effects of AA, AA3, and myxothiazol on PSII activity and linear photosynthetic electron transport using isolated plant PSII and thylakoid membranes. AA, but neither AA3 nor myxothiazol, directly suppressed PSII activity and linear electron transport. Furthermore, the extent of AA’s effects was batch-dependent. Thus, we propose using AA3 to inhibit CEF and myxothiazol to inhibit complex III, instead of AA.

**Graphical Abstract:** 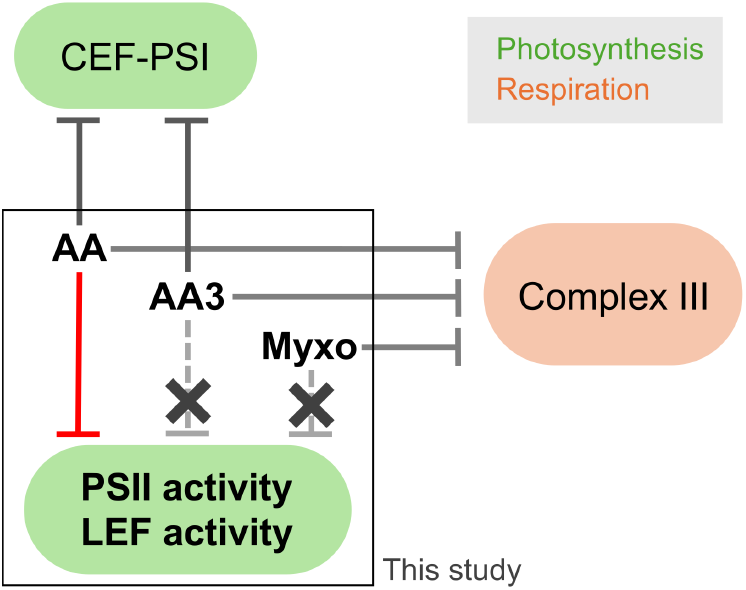

Antimycin A inhibits at least three sites. Antimycin A3 has higher specificity for CEF-PSI, while myxothiazol is preferred to inhibit complex III.

---

Photosynthetic electron transport consists of both linear electron flow (LEF) and cyclic electron flow around photosystem I (CEF-PSI). LEF is initiated at photosystem II (PSII), the supercomplex that uses light energy to oxidize water into molecular oxygen at its donor side, and reduces plastoquinone (PQ) at its acceptor side (Imaizumi and Ifuku, 2025). The electrons are then sequentially transferred from PSII through the PQ pool, cytochrome (Cyt) *b*_6_*f*, plastocyanin (or Cyt *c*_6_), PSI, and ferredoxin (Fd), and are finally used to reduce NADP^+^ to NADPH via Fd-NADP^+^ reductase (FNR) (Johnson, 2025). On the other hand, CEF-PSI does not incorporate PSII, and electrons are transferred from reduced Fd back to the PQ pool. One CEF-PSI pathway is mediated by the NADH dehydrogenase-like (NDH) complex and is known as the NDH-dependent CEF-PSI pathway (Ifuku et al., 2011; Shikanai et al., 2025). Another proposed pathway, which has been suggested to be the major CEF-PSI pathway, relies on the protein PROTON GRADIENT REGULATION 5 (PGR5) and is thus referred to as the PGR5-dependent CEF-PSI pathway (Munekage et al., 2002). Extensive studies have revealed the importance of PGR5 (Yamori and Shikanai, 2016); the molecular mechanism of this pathway, however, remains elusive (Johnson, 2025).

In the field of photosynthesis, antimycin A (AA), has been known as an inhibitor of CEF-PSI for over half a century (Tagawa et al., 1963). Studies have suggested that AA inhibits PGR5-dependent CEF-PSI but not NDH-dependent CEF-PSI (Joët et al., 2001; Munekage et al., 2002, 2004), yet the mechanism by which AA acts remains poorly understood. Nevertheless, numerous studies have since used AA to inhibit the PGR5-dependent pathway (Labs et al., 2016), often regarding it as a specific inhibitor. Meanwhile, there have been several studies suggesting that AA can also affect PSII (Hind, 1968; Cramer et al., 1971; Cramer and Böhme, 1972; Satoh and Katoh, 1972; Yerkes and Crofts, 1992; Miyake et al., 1995; Takagi et al., 2019), most of them revealing that it alters the redox properties of Cyt *b*_559_, which is part of the core of PSII. However, these reports have often been overlooked.

AA is also known to inhibit complex III (Cyt *bc*_1_) of the mitochondrial respiratory chain (Slater, 1973) by binding to the Q_i_ site. Myxothiazol is another well-known inhibitor of complex III (Thierbach and Reichenbach, 1981; von Jagow and Engel, 1981) which binds to the Q_o_ site. As AA inhibits both complex III and the photosynthetic CEF-PSI, myxothiazol is considered to be a more specific inhibitor than AA, in terms of inhibiting complex III. However, it is not well-studied whether myxothiazol does not have any direct effects on photosynthetic electron transfer.

Recently, we found that AA has remarkable direct inhibitory effects on PSII in the presence of Q_B_-site binding inhibitors such as DCMU (Imaizumi et al., 2025). While AA by itself also showed direct inhibitory effects on electron transfer within PSII (Imaizumi et al., 2025), it remains unclear to what extent AA can affect PSII or LEF in the absence of Q_B_-site binding inhibitors. We also observed that myxothiazol can have some direct effect on PSII (Imaizumi et al., 2025), but the extent of its impact on PSII or LEF activity remains elusive, too. Conventionally used AA is a mixture of closely related compounds AA1, AA2, AA3, and AA4 (Liu and Strong, 1959). We discovered that AA1 and AA2 have inhibitory effects on PSII, whereas AA3 and AA4 do not (Imaizumi et al., 2025). Since AA3 efficiently inhibited PGR5-dependent CEF-PSI as AA does (Imaizumi et al., 2025), consistent with a previous study (Taira et al., 2013), the use of AA3 enables inhibition of the PGR5-dependent pathway without directly affecting PSII. In this study, we investigated the direct impacts of AA, AA3, and myxothiazol (**Figure 1**) on the whole-chain electron transport activity in photosynthesis and on the oxygen-evolving activity of PSII. The results provide insight into improved ways of using these chemical inhibitors to study photosynthesis and respiration in photosynthetic organisms.

**Figure 1.**
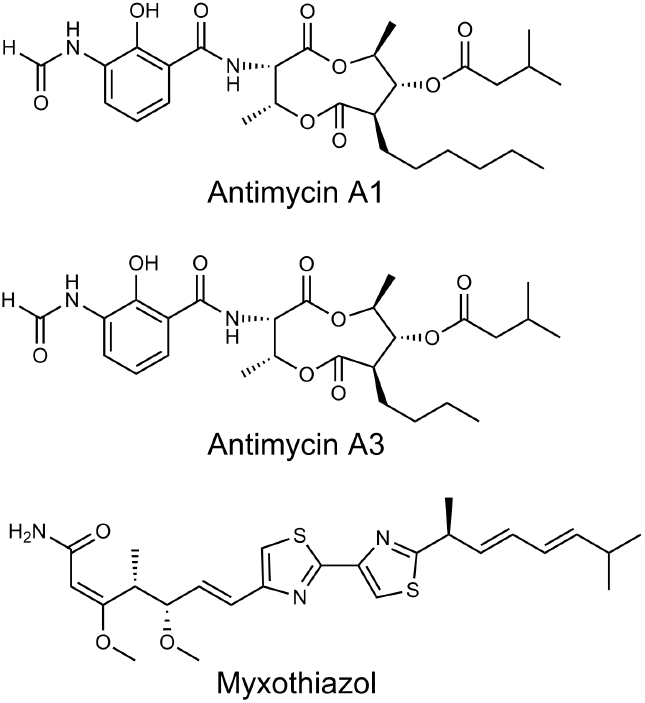
Chemical structures of AA1, AA3, and myxothiazol.

## Materials and methods

### Growth conditions of *Arabidopsis thaliana*

Wild-type (Col-0; Columbia-0) and PGR5-deficient mutant (*pgr5*^*hope1*^) *A. thaliana* were grown for one week on half-strength Murashige and Skoog (MS) medium solidified with 0.9% (w/v) agar. Then, seedlings were transferred to soil and grown for two more weeks. Leaves from three-week-old plants were used for the isolation of *Arabidopsis* thylakoid membranes. Plants were grown at 22 ºC under long-day conditions (16 h light/8 h dark) at a light intensity of 80 µmol photons m^−2^ s^−1^.

### Isolation of PSII membranes and thylakoid membranes

Oxygen-evolving PSII membranes, known as BBY membranes (Berthold et al., 1981), were isolated from market spinach based on Yamamoto et al. (Yamamoto et al., 2011) as reported previously (Imaizumi et al., 2022); the oxygen-evolving activity in the absence of inhibitors was between 370 and 520 µmol O_2_ mg Chl^−1^ h^−1^. Isolation of spinach thylakoid membranes was conducted by modifying this method as reported previously (Imaizumi et al., 2025); the whole-chain electron transport activity (oxygen uptake activity) in the absence of inhibitors was between 205 and 262 µmol O_2_ mg Chl^−1^ h^−1^. Isolation of *A. thaliana* thylakoid membranes was conducted by slightly modifying the methods by Allahverdiyeva et al. (2007) as reported previously (Imaizumi et al., 2025); the whole-chain electron transport activity (oxygen uptake activity) of Col-0 thylakoids in the absence of inhibitors was approximately 120 µmol O_2_ mg Chl^−1^ h^−1^.

### Chemical inhibitors

AA (mixture of AAs isolated from *Streptomyces* sp.) was purchased from Sigma-Aldrich (product number A8674). Unless indicated, we used AA (lot number 125K4030), whose major components were AA1 (73%), AA2 (19%), AA3 (5%), and AA4 (1%) according to the Certificate of Analysis by Sigma-Aldrich. When comparing different batches of AA with the same product number, we also used AA (lot number 0000187236), which will hereafter be referred to as “AA (mix#2)”, whose major components were AA1 (34%), AA2 (17%), AA3 (34%), and AA4 (13%) according to the Certificate of Analysis by Sigma-Aldrich. AA1 was purchased from BioAustralis Fine Chemicals (product number BIA-A1442; batch AC33.74). AA3 was purchased from BioAustralis Fine Chemicals (product number BIA-A1444; batch AC32.10). Myxothiazol was purchased from Sigma-Aldrich (product number T5580; batch 0000189591). All inhibitor stock solutions were prepared in ethanol.

### Measurement of PSII activity and whole-chain electron transport activity

Oxygen-evolving activity of PSII (10 µg Chl mL^−1^) was measured using a Clark-type oxygen electrode at 25 ºC in buffer A (20 mM HEPES-NaOH, 1 M betaine, 10 mM NaCl, 5 mM MgCl_2_, and pH 7.6) with 0.5 mM phenyl-*p*-benzoquinone (PPBQ) as electron acceptor under saturating light illumination (2,500 µmol photons m^−2^ s^−1^). Whole-chain electron transport activity of thylakoid membranes (10 µg Chl mL^−1^) was examined by measuring the oxygen uptake activity using a Clark-type oxygen electrode at 25 ºC in buffer A with 100 µM methyl viologen (MV) as electron acceptor in the presence of 1 mM KCN, and 5 mM NH_4_Cl as uncoupler under saturating light illumination.

## Results

### Antimycin A, but not antimycin A3, suppresses linear electron flow

Many studies have shown that AA affects photosynthetic electron transport (Labs et al., 2016). While these effects have often been attributed to the inhibition of PGR5-dependent CEF-PSI or the inhibition of complex III in the mitochondrial respiratory chain, there may also be other direct inhibitory effects of AA (Yerkes and Crofts, 1992; Fisher and Kramer, 2014; Takagi et al., 2019; Buchert et al., 2022; Imaizumi et al., 2025). Therefore, we first investigated the effects of AA on the whole-chain electron transport activity in photosynthesis. To eliminate any effects of inhibition of mitochondrial respiration, we used isolated spinach thylakoid membranes. AA was used at 10 µM, which is a commonly used concentration for inhibition of CEF. The AA we used was composed of AA1 (73%), AA2 (19%), AA3 (5%), and AA4 (1%). AA significantly suppressed the whole-chain activity to approximately 70–80% of the control (**Figure 2**). Similarly, AA1, which is the AA component with notable effects on PSII (Imaizumi et al., 2025), also suppressed the whole-chain electron transport activity. Meanwhile, AA3, which inhibits PGR5-dependent CEF-PSI without affecting PSII (Imaizumi et al., 2025), did not affect the whole-chain activity. These results implied that the observed suppression of whole-chain activity by AA may be due to its inhibitory effects on PSII, but not due to inhibition of PGR5 functions. Similar effects of AA on the whole-chain activity were also observed using thylakoid membranes isolated from Col-0 (wild-type) and *pgr5*^*hope1*^ (PGR5-deficient mutant) *Arabidopsis thaliana*, confirming that the observed AA effects are distinct from the effects of AA on PGR5 functions (**Figure 3**).

**Figure 2.**
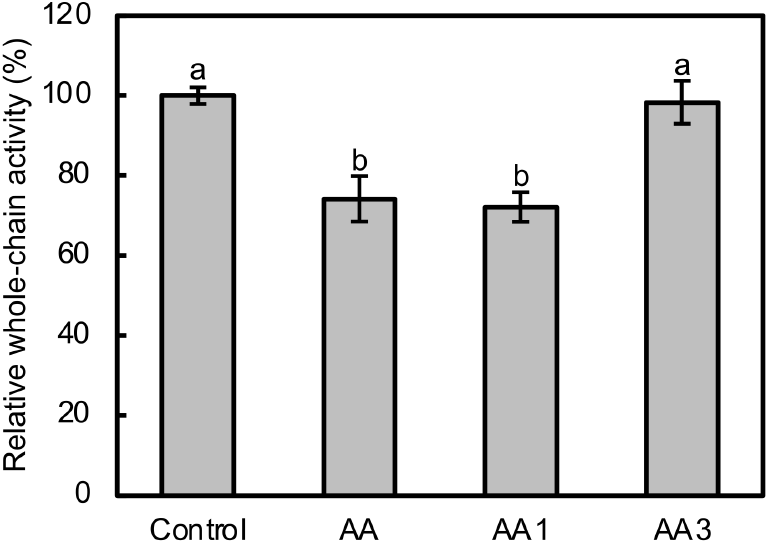
Direct effects of AA, AA1, and AA3 on the whole-chain electron transport activity of isolated spinach thylakoid membranes. Inhibitors were added to a final concentration of 10 µM and whole-chain (water to MV) activity was measured in the presence of 100 µM MV as electron acceptor. Data are mean ± SD (*n* = 4), and activity without inhibitors was set as 100%. Different lowercase letters above bars indicate a statistically significant difference (*P* < 0.05, Tukey’s HSD test).

**Figure 3.**
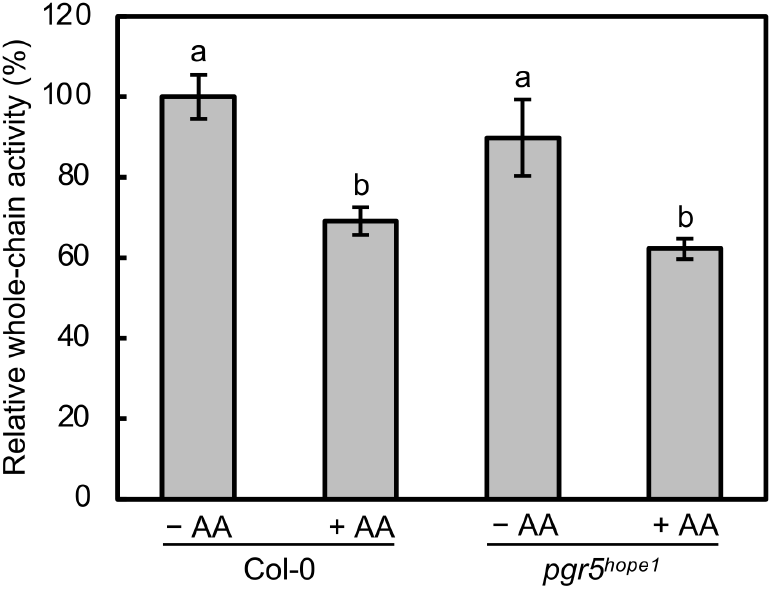
Direct effects of AA (10 µM) on the whole-chain electron transport activity of *Arabidopsis* thylakoid membranes isolated from Col-0 and *pgr5*^*hope1*^. Whole-chain activity was measured in the presence of 100 µM MV as electron acceptor. Data are mean ± SD (*n* = 3), and activity of Col-0 without inhibitors was set as 100%. Different lowercase letters above bars indicate a statistically significant difference (*P* < 0.05, Tukey’s HSD test).

### Antimycin A, but not antimycin A3, directly suppresses PSII activity

To clarify whether AA directly suppresses PSII activity, we next investigated the effects of AA on the oxygen-evolving activity of PSII using isolated spinach PSII membranes (**Figure 4**). AA significantly suppressed the oxygen-evolving activity of PSII to approximately 80% of the control. This confirms that AA indeed has direct inhibitory effects on PSII activity. Meanwhile, AA3 did not have any effects on the oxygen-evolving activity of PSII. This is consistent with our previous study, which showed that AA has inhibitory effects on PSII, while AA3 does not affect PSII (Imaizumi et al., 2025). These results strongly suggest that the inhibitory effects of AA are not restricted to PGR5-dependent CEF-PSI and complex III of the mitochondrial respiratory chain; it can have significant impacts on PSII.

**Figure 4.**
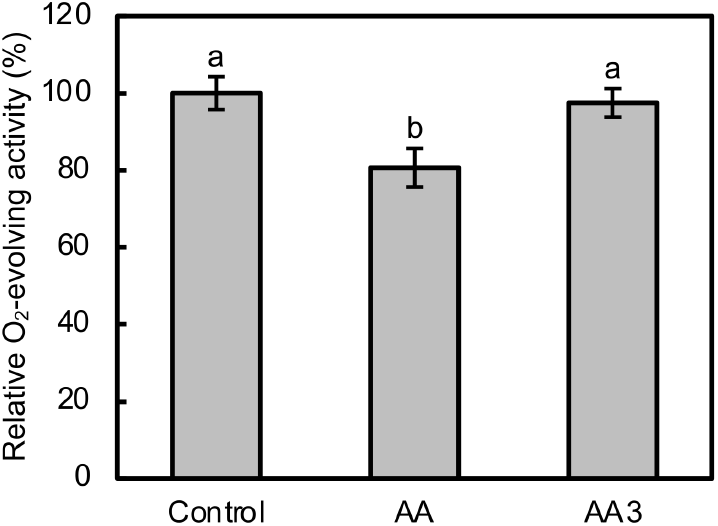
Direct effects of AA (10 µM) and AA3 (10 µM) on the oxygen-evolving activity of isolated spinach PSII membranes. Oxygen-evolving activity was measured in the presence of 0.5 mM PPBQ as electron acceptor. Data are mean ± SD (*n* = 3), and activity without inhibitors was set as 100%. Different lowercase letters above bars indicate a statistically significant difference (*P* < 0.05, Tukey’s HSD test).

### Myxothiazol does not directly affect linear electron flow significantly

AA and myxothiazol are well-known inhibitors of complex III. In contrast to AA, known to also inhibit PGR5-dependent CEF-PSI, myxothiazol is often thought to be a more specific inhibitor of complex III. However, few studies have clarified whether myxothiazol has direct effects on photosynthetic electron transport or not. Therefore, we also examined the effects of myxothiazol on the whole-chain electron transport activity of isolated thylakoid membranes and on the oxygen-evolving activity of isolated PSII membranes. Myxothiazol did not have significant effects on the whole-chain activity (**Figure 5a**). We also did not observe significant effects of myxothiazol on the oxygen-evolving activity of isolated PSII membranes (**Figure 5b**). These results confirm that myxothiazol is indeed a more specific inhibitor than AA both in terms of their effects on photosynthetic LEF and on photosynthetic CEF-PSI.

**Figure 5.**
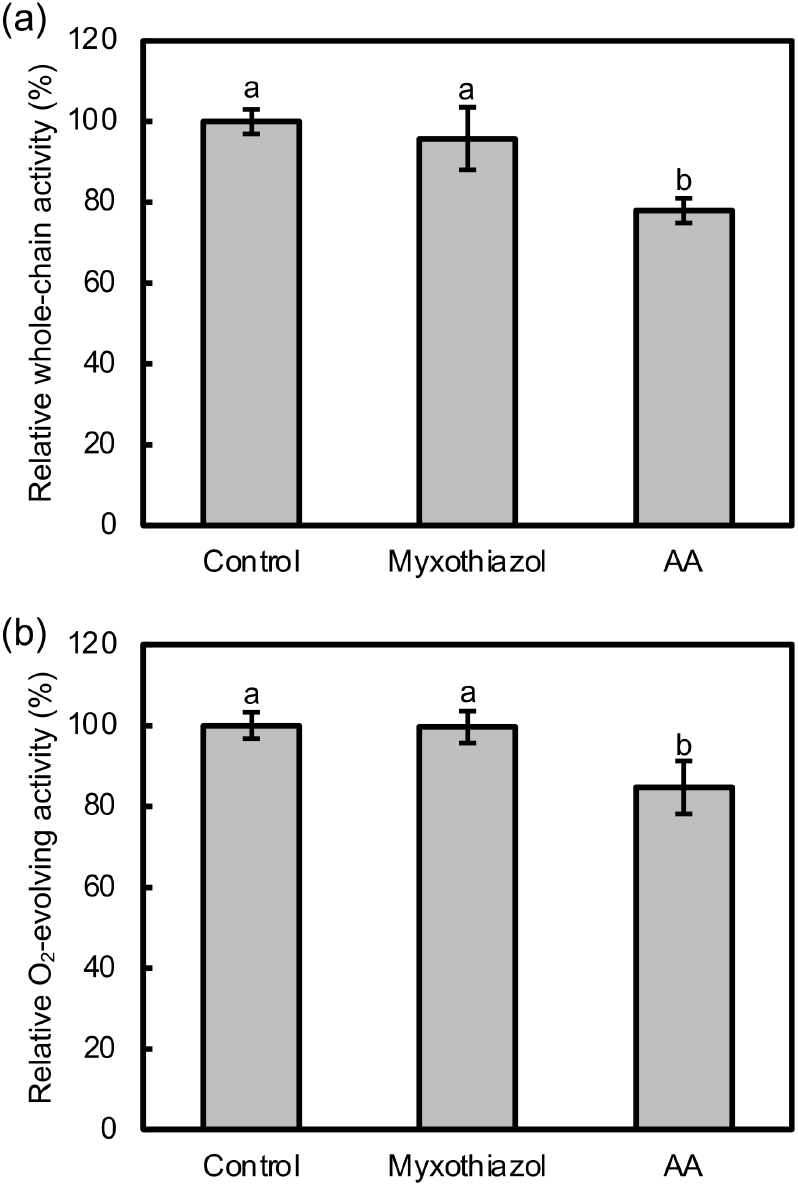
Direct effects of myxothiazol (10 µM) on photosynthetic electron transport. (a) Whole-chain activity (*n* = 3) of isolated spinach thylakoid membranes was measured in the presence of 100 µM MV as electron acceptor, and (b) oxygen-evolving activity (*n* = 4) of isolated spinach PSII membranes was measured in the presence of 0.5 mM PPBQ as electron acceptor. Data are mean ± SD and activity without inhibitors was set as 100%. Different lowercase letters above bars indicate a statistically significant difference (*P* < 0.05, Tukey’s HSD test).

## Discussion

AA is commonly used for the inhibition of PGR5-dependent CEF-PSI. Many studies on photosynthetic electron transport have used AA as if it were a specific inhibitor of PGR5-dependent CEF-PSI, without considering the possibility of its direct effects on PSII or LEF. Our findings demonstrate that AA can actually have direct and significant inhibitory effects on the activity of PSII, which does not participate in CEF.

The whole-chain electron transport activity of isolated thylakoid membranes was notably suppressed by AA (or AA1), but not by AA3 (**Figure 2**). The suppression was also observed using thylakoid membranes isolated from *pgr5*^*hope1*^ mutant *A. thaliana* (**Figure 3**). These results clarify that AA has direct effects on LEF other than its effects on CEF or on the mitochondrial respiratory chain. We have recently shown that of the four major AA components, AA1 and AA2 have inhibitory effects on PSII, whereas AA3 and AA4 do not (Imaizumi et al., 2025). Therefore, the observation that AA and AA1 but not AA3 suppressed the whole-chain activity suggested that the site of suppression may be PSII. This has been confirmed by measurements using isolated PSII membranes (**Figure 4** and **Figure 5b**). We do not rule out the possibility that AA could also have direct effects on Cyt *b*_6_*f* (Buchert et al., 2022). However, AA1 and AA3 are structurally very similar (**Figure 1**), and both of them inhibit PGR5-dependent CEF-PSI (Imaizumi et al., 2025) as well as complex III; the case with PSII, with only one of them having inhibitory effects (Imaizumi et al., 2025), is more likely to be a rare case. Thus, the suppression of whole-chain activity by AA that we observed seems to be due to its effect on PSII. It should be noted that the relative proportions of AA1 to AA4 vary considerably between different batches of AA (Imaizumi et al., 2025), meaning that the effects of AA on photosynthesis can differ depending on the composition of the batch that is used. In fact, using two different batches of AA with the same product number, we confirmed that the effects of AA on the whole-chain activity depends on the ratio of the AA components (**Figure 6**). While AA with high AA1 content (73%) significantly suppressed the activity to approximately 80%, a different batch of AA (AA [mix#2]) containing similar proportions of AA1 (34%) and AA3 (34%) only showed a tendency to suppress the activity to approximately 90%. Therefore, the use of AA is not recommended, but whenever AA is used, at a minimum, its batch number and composition should be reported.

**Figure 6.**
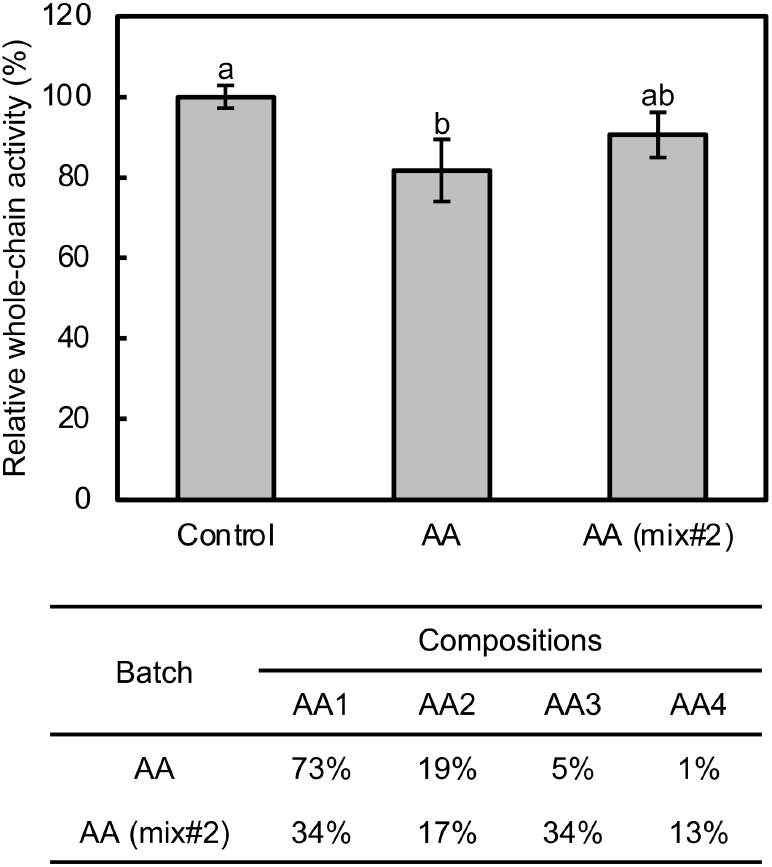
Comparison of the direct effects of two different batches of AA (10 µM) on the whole-chain electron transport activity of isolated spinach thylakoid membranes. Whole-chain activity was measured in the presence of 100 µM MV as electron acceptor. The two batches of AA (“AA” and “AA [mix#2]”) showed considerable differences in the proportions of AA1 to AA4. Data are mean ± SD (*n* = 3), and activity without inhibitors was set as 100%. Different lowercase letters above bars indicate a statistically significant difference (*P* < 0.05, Tukey’s HSD test).

AA also suppressed the oxygen-evolving activity of PSII in isolated PSII membranes (**Figure 4**). This observation provides clear evidence that AA can have direct inhibitory effects on PSII activity. The inhibitor concentration (10 µM) we used throughout this study is commonly used for inhibiting CEF-PSI. Moreover, we have already studied the concentration dependence of AA effects on PSII previously, and showed that the inhibitory effects of AA on PSII reaches near-saturation at this concentration (Imaizumi et al., 2025). Considering that various Q_B_-site binding inhibitors and AA can bind to PSII simultaneously, the binding site of AA is most likely not the Q_B_ site. Instead, AA has been suggested to bind near the acceptor side of PSII (e.g., in a PQ exchange channel) to modulate the local environment, leading to decreased efficiency, but not full inhibition, of electron transfer from Q_A−_ to Q_B_ and of the exchange of PQ molecules (Imaizumi et al., 2025). Therefore, the current results showing that PSII activity is suppressed to approximately 80% in the presence of sufficient AA are consistent with our previous findings.

While our recent report revealed remarkable impacts of AA on PSII under the presence of Q_B_-site binding inhibitors such as DCMU (Imaizumi et al., 2025), the current study adds to it, showing that there are also significant impacts in the absence of additional inhibitors other than AA. Therefore, when AA is used as an inhibitor of CEF, the results can contain consequences of undesired effects of AA on PSII, as well. Incorrect assumptions that AA is a CEF-specific inhibitor have led to conclusions drawn without adequate consideration of alternative explanations. For example, many reports have shown decreased PSII activity and/or increased PSII photoinhibition by AA treatment, and these effects have often been attributed to inhibition of CEF. It is possible that some of these effects were at least partially due to direct effects of AA on PSII. The fact that CEF is often studied by measuring the chlorophyll fluorescence that derive from PSII may make the presence of direct AA effects on PSII even more problematic.

Although AA is also used as an inhibitor of complex III of the mitochondrial respiratory chain, due to the widely recognized effects of AA on PGR5-dependent CEF-PSI, myxothiazol is often believed to be a more specific alternative inhibitor of complex III. Meanwhile, the direct effects of myxothiazol on the photosynthetic electron transport chain are not thoroughly studied, and in fact, we previously observed that it can have some direct effects on PSII (Imaizumi et al., 2025). This has driven us to confirm whether or not myxothiazol affects PSII and/or LEF activity. As a result, we did not observe significant effects of myxothiazol in the current assays, in contrast to AA having remarkable effects (**Figure 5**). Complex III has been suggested to be more sensitive to AA than photosynthetic electron transport pathways are, and therefore, using AA at low concentrations could be effective in increasing its specificity to mainly inhibit respiration (Watanabe et al., 2016). However, considering that low concentrations of AA (1 µM or less) could also affect PSII (Imaizumi et al., 2025) and CEF (Sugimoto et al., 2013; Taira et al., 2013) to some extent, the use of myxothiazol would still be preferential.

## Conclusion

Chemical inhibitors are powerful tools for dissecting biological pathways, but improper use or incorrect assumptions about their specificity can result in misleading interpretations. AA has often been misunderstood as a specific inhibitor of PGR5-dependent CEF-PSI in the photosynthetic electron transport pathways. However, we have shown that AA has direct inhibitory effects on PSII and cannot be used as a specific inhibitor, even within the chloroplast. Meanwhile, AA3, a commercially available component of AA, inhibits PGR5 functions as AA does, but does not show direct effects on PSII. It should be noted that AA3 most likely also inhibits plant and algal complex III as AA does (Liu and Strong, 1959). In terms of inhibiting complex III of the mitochondrial respiratory chain, myxothiazol did not show direct significant effects on PSII or on LEF and is most likely more specific to complex III compared to AA. Thus, to chemically inhibit PGR5-dependent CEF-PSI, AA3 should be used instead of AA (with myxothiazol serving as control, if necessary), and to inhibit complex III, myxothiazol should be used instead of AA.

## Data Availability

The data underlying this article are available in the article and its online supplementary data.

## Author Contributions

K.I. and K.If. conceived the project; K.I. performed all the analyses and drafted the original manuscript; K.I. and K.If. revised the manuscript and prepared the final manuscript.

